# Mechanical loading of cranial joints minimizes the craniofacial phenotype in Crouzon syndrome

**DOI:** 10.1101/2021.08.31.458360

**Authors:** Mehran Moazen, Mahbubeh Hejazi, Dawn Savery, Dominic Jones, Arsalan Marghoub, Ali Alazmani, Erwin Pauws

**Author notes:** These authors contributed equally (Mehran Moazen, Mahbubeh Hejazi).

## Abstract

Children with syndromic forms of craniosynostosis undergo a plethora of surgical interventions to resolve the clinical features caused by the premature fusion of cranial sutures. While surgical correction is reliable, the need for repeated rounds of invasive treatment puts a heavy burden on the child and their family.

This study explores a non-surgical alternative using mechanical loading of the cranial joints to prevent or delay craniofacial phenotypes associated with Crouzon syndrome.

We treated Crouzon syndrome mice before the onset of craniosynostosis by cyclical mechanical loading of cranial joints using a custom designed set-up.

Cranial loading applied to the frontal bone partially restores normal skull morphology, significantly reducing the typical brachycephalic appearance. This is underpinned by the delayed closure of the coronal suture and of the intersphenoidal synchondrosis.

This study provides a novel treatment alternative for syndromic craniosynostosis which has the potential to be an important step towards replacing, reducing or refining the surgical treatment of all craniosynostosis patients.

## 1. Introduction

Some of the most common birth defects affect the craniofacial development of a new-born baby. Therefore, the care of children with craniofacial birth defects is an important clinical activity. Diagnosis and treatment of the disorders typically involves a child being seen by a number of specialists in areas including clinical genetics, radiology, dentistry, maxillo-facial and neurosurgery, and anaesthesia. Craniosynostosis is a common birth defect with an estimated prevalence between 1 in 2,100^1^ and 1 in 4,200 incidence^2^. It is caused by the premature fusion of the bones that form the skull, which can have serious, possibly life-threatening results^3,4^. Traditional approaches to the treatment of this type of craniofacial birth defect employ complex surgical remodelling of the skull including facial deformities, aimed at protecting brain development, visual function and the restoration of a more normal craniofacial appearance.

The most severe forms of craniosynostosis that require the most intensive treatments are often syndromic and single-gene disorders^5^. Surgical treatment of syndromic craniosynostosis can be clinically challenging because in many cases multiple sutures are affected. Also, the genetic and phenotypic variability between patients, even those within the same syndrome classification is often significant enough to require individualised treatment protocols^6,7^. In addition, these patients require an increased number of craniofacial procedures^5^. The number of post-surgical complications is higher in syndromic cases, with increased rates seen in secondary synostosis^8^. There is also a high number of complication associated with frontonasal distraction procedures^9^.

While the underlying pathogenesis of syndromic cases of craniosynostosis is complex and can involve a plethora of developmental signalling pathways, the majority of cases are caused by mutations in genes directly or indirectly involved in the FGF signalling pathway^10^. In addition, abberant FGF signalling plays an important role in many other developmental skeletal defects, affecting endochondral as well as intramembranous ossification^11^. Besides abnormalities in cellular physiology, studies showing an association between intrauterine constraint and increased incidence of craniosynostosis indicate a role for external mechanical forces on the formation and/or homeostasis of cranial sutures^12^.

Despite the rapid evolution of human genetics, gene therapy and stem cell therapy during the last few decades, there are currently no suitable alternatives to surgery. While our understanding of the pathogenesis of syndromic craniosynostosis has increased significantly -mainly through the use of genetically manipulated mouse models-this has not resulted in any clinically applicable non-surgical treatment alternatives^13^.

Mechanotransduction is the process whereby cells convert mechanical stimulus into a molecular and cellular response. While there is a large body of literature on mechanical loading of endochondral bone^14^, only recently studies have started to explore how intramembranous, craniofacial bones and sutures respond to mechanical forces. These studies have shown that craniofacial sutures respond to cyclical, mechanical loading using both in vitro and *in vivo* models. Resected calvaria of wild-type rat skulls were loaded on the posterior intrafrontal suture under tension in an ex vivo model. It was reported that loaded sutures remained patent while in the control group the suture fused normally^15^. *In vivo* cyclical loading applied to the skull (i.e. maxilla) has been shown to increase the width of the premaxillomaxillary suture by approximately 30% compared to controls in rat^16^ and similar results have been obtained in rabbit^17^. Normal craniofacial sutures in a pig model have also been shown to respond to tension forces by proliferation and differentiation of suture cells, while compression forces lead to increased osteogenesis^18^. Therefore, the aforementioned studies suggest that cyclical, mechanical loading can keep cranial sutures patent during postnatal development and delay their natural closure.

Many mouse models have been developed that feature the craniosynostosis phenotype^13^. For this study we used an established model of Crouzon syndrome. Crouzon mice carry a mutation (p.C342Y) in the FGFR2 receptor, are viable and fertile and are characterised by brachycephaly caused by coronal craniosynostosis as well as midfacial hypoplasia associated with malocclusion^19^. Our laboratory has been studying this mouse model since 2010 and have built up considerable knowledge of the cellular and molecular events underlying the pathogenesis of the phenotypic features this animal model exhibits, including impaired mesenchymal condensation of skeletal elements due to misregulation of SOX9^20^. A study from Hatch et al. showed that genetic background effects phenotypic severity with a more pronounced craniofacial phenotype in C57BL/6 mice compared to BALB/c^21^. Our mice have been backcrossed onto the CD-1 strain, which mimicks human Crouzon syndrome and in our hands displays synostosis of the coronal suture at three weeks of age (P21). We have previously shown differences in biomechanical properties of individual calvarial bones (i.e. frontal bone vs parietal bone) in Crouzon syndrome calvarial bones and sutures^22^. These observations point to a fundamental difference between the architecture of normal craniofacial bone and suture tissue affected by FGFR2 mutation. These observations are supported by studies showing that the neural crest derived frontal bone is more susceptive to FGFR2 perturbation than the mesoderm derived parietal bone^23,24^.

The aim of this study was to investigate the possibility that external mechanical force applied to cranial bone can delay or prevent closure of abnormal sutures prone to craniosynostosis. Previous research has shown that mechanical loading of cranial bone in can delay the normal closure of cranial sutures during postnatal development. Our hypothesis was that *in vivo* mechanical loading of calvarial bone can significantly delay premature, pathogenic suture closure in a mouse model for craniosynostosis.

## 2. Materials and methods

### 2.1. Animals

#### Crouzon mouse model (*Fgfr2*^tm4Lni^; aka *Fgfr2c*^C342Y^; MGI:3053095)

*Fgfr2c*^C342Y^ were re-derived through the European Mouse Mutant Archive (EMMA) at MRC Harwell as previously described^20^.

All animal experiments were approved by the UK Home Office and performed as part of a Project License (number: 70/8817) under the UK Animals (Scientific Procedures) Act 1986. Animal procedures procedures complied with the ARRIVE guidelines and were performed under the supervision of UCL Biological Services.

### 2.2. Mechanical loading

An experimental loading set up was developed to precisely load the cranial bones of mice *in vivo*. Here an actuator (T-LSR series, Zaber Technologies: res. 50 μm, max. load 200 N) and a force sensor (GSO Series, Transducer Techniques: res. 0.01 N with 1N capacity) were configured to a custom developed LabVIEW program (National Instruments Corp, Austin, TX, USA). This portable set up enabled us to carry out precise cyclic loading of mouse skulls.

### 2.3. Phenotypic analysis

Animals were weighed using a fine balance. An electronic caliper (Fisher) was used to measure the crown-rump length and head length of the animal.

### 2.4. Micro-CT analysis

#### 2.4.1. Micro-CT scanning

At the end of experiment, animals were killed by a schedule 1 method or a non-schedule 1 method (asphyxiated using rising concentrations of CO2). Then animals were fixed and stained with Alizarin Red to better visualise the calvarial bones and sutures. They were then micro CT scanned (X-Tek HMX 160, XTek Systems Ltd, Tring, Herts., UK).

#### 2.4.2. Image processing

CT images were imported into an image processing software (Avizo Lite, FEI V9.2., Thermo Fisher Scientific, Mass, USA). The quality of micro-CT images, voxel sizes, were between 14-20μm. Here the bone was segmented. Before segmentation micro-CT images were aligned in a similar position in different planes. Following the alignment of the raw CT images, 3D surface volumes were reconstructed and a range of parameters were measured.

#### 2.4.3. Quantitative analysis

In the lateral view, the differences in the length, height and the overall morphology of the calvarial were observed between the groups. The dorsal view, highlights the suture fusion across the calvaria across the groups. The skull length was measured in the sagittal plane between the most anterior-medial point of the nasal bone and the most posterior-medial point of the occipital bone. Skull width was then measured in the transverse plane between the most posterior-inferior point on the interparietal. Skull height was measured in the sagittal plane between the presphenoid and the most superior part of the skull, covering bulging area in the mutant types. The aforementioned measurements were normalised by calculating the relative differences between the experimental groups and the control group (wild-type untreated) and presenting these as a percentage increase/decrease.

#### 2.4.4. Patency scoring of synchondroses

Sagittal sections through the midline of whole skull micro-CT images were presented to at least three observers blinded for treated and untreated groups. SOS and SIS synchondroses were scored as open, partially closed (<50% of the cranial base height is filled with bone) or closed (<50% of the cranial base height is filled with bone). Scores were averaged and plotted against expected frequencies in untreated groups.

### 2.5. Fluorescent Alizarin Red staining

Mice were injected intraperitoneally with 5ul/gram body weight of a 5mg/ml Alizarin complexone (VWR 20118.081) made up in 2% NaHCO3 in PBS. Calvaria were dissected 24 hours after injection and imaged using a Zeiss upright fluorescent microscope with a 550nm filter.

### 2.6. Histological analysis

For paraffin embedding, mouse heads were skinned and fixed in 10% Formalin overnight before graded dehydration in ethanol. Embryos were cleared in analytical grade xylene (Fisher) before paraffin wax displacement in a 60°C oven. The samples were embedded and sectioned between 8-10um on the sagittal plane using a microtome (Leica). Sections were stained with Haematoxylin and Eosin.

### 2.7. Statistical analysis

SPSS Statistics 22 (IBM) software was used as the primary statistical package for data analysis. First, the data was tested for normality using Shapiro-Wilk test to determine the use of parametric or non-parametric tests. Independent samples T-test with Welch’s correction was used to compare the difference of averages between groups for quantification. Chi^2^-test was used for statistical analysis of the synchodrosis patency score. One-way ANOVA with Tukey posthoc or non-parametric Kruskal Wallis with Dunn-Bonferroni posthoc test was adopted for analysis of three or more groups. A p-value of <0.05 was considered significant.

### 2.8. Finite element analysis

A previously validated finite element model of mouse skull growth was used here^25,26^. In brief, we reconstructed micro-CT images of a P3 WT mouse (Avizo Lite, FEI V9.2., Thermo Fisher Scientific, Mass, USA) consisted of bone, sutures, and intracranial volume (ICV) that broadly represented the brain. The whole model was then transformed into a 3D solid meshed model and imported to a finite element solver, ANSYS v.18 (ANSYS Inc., Canonsburg, PA, USA). Isotropic (linear and elastic) material properties were assigned to all sections with a thermal coefficient defined only for the ICV. Bone and suture were assumed to have an elastic modulus of 3500 MPa and 30 MPa respectively at age P3^22,27^. The elastic modulus of the ICV was assumed to be 10 MPa. The bone and suture materials were assumed to have a Poisson’s ratio of 0.3. The ICV value was 0.48. The bone-suture interfaces and bone-suture-intracranial volume interfaces were assumed to be perfectly connected. Three nodes on the presphenoid bone were constrained in all degrees of freedom. The presphenoid bone was constrained based on our previous examination of the growth of the WT mouse skull that revealed that this bone grows centrically during development and can be considered to effectively remain at the same position during the skull development^25^. The P3 model was then virtually grown to P7 (as described in Marghoub et al.^26^) and at P7 was loaded at the fronal and parietal analogoun to our *in vivo* experiment. The pattern of first principal strain across the skull were then compared between the two groups.

## 3. Results

### 3.1. *In vivo* mechanical loading of cranial bone in a Crouzon syndrome mouse model

To test our hypothesis that mechanical loading of the skull can prevent pathogenic suture closure, we performed *in vivo* loading of cranial bones on Crouzon mice. Using pre-weaning mice allowed us to use both male and female animals in order to compare littermates with sufficient numbers (n≥3). We developed a bespoke system using a linear actuator and force sensor enabling us to carry out controlled and repeatable loading of the cranial bones (**Figure 1a**). The actuator moved a force sensor in the dorso-ventral direction while a probe was fixed to it, in effect loading the skull via the probe. The loading tip of the probe that came into contact with the skull had a flat circular surface with a diameter of 1 mm. Based on previous work on rabbit and rat models ^16,17^, we decided to apply a force of 10 gram at 1Hz for all our cranial bone loading protocols. To facilitate accurate, positional loading on cranial bones of pre-weaning mice, we used general aneasthesia by Isoflurane inhalation to fix animals in position using a custom-made, 3D printed tube underneath the actuator probe for the duration of the loading procedure (**Figures 1B and 1C**). We loaded all animals on the left side of the cranium to allow us to compare with the right side as an unloaded control (**Figure 1D**). Because of the overlapping nature of the coronal suture (**Figure 1E**) we decided to perform loading experiments on the frontal and parietal bone seperately. This way, we were able to compare the application of tension force on the suture when loading on the frontal bone and compression force on the suture when loading on the parietal bone (**Figure 1F**). We used an Fgfr2-C342Y transgenic mouse line that is kept on a CD-1 background, that in our hands displays patent sutures at birth, followed by the onset of craniosynostosis which is usually complete by P21. To maximise the potential impact of our cranial bone loading treatment, we loaded animals for 10 days between P7-18. Skulls were subsequently dissected and analysed at P21 (**Figure 1G**). **Supplementary Videos 1a and 1b** show examples of the cranial bone loading procedure at P7 and P14 respectively. In all experiments, mice recovered quickly after daily cycles of aneasthesia and cranial bone loading and at P21, after 10 rounds of treatment, animals did not show signs of abnormal growth or behaviour (**Supplementary Video 2a and b**).

**Figure 1.**
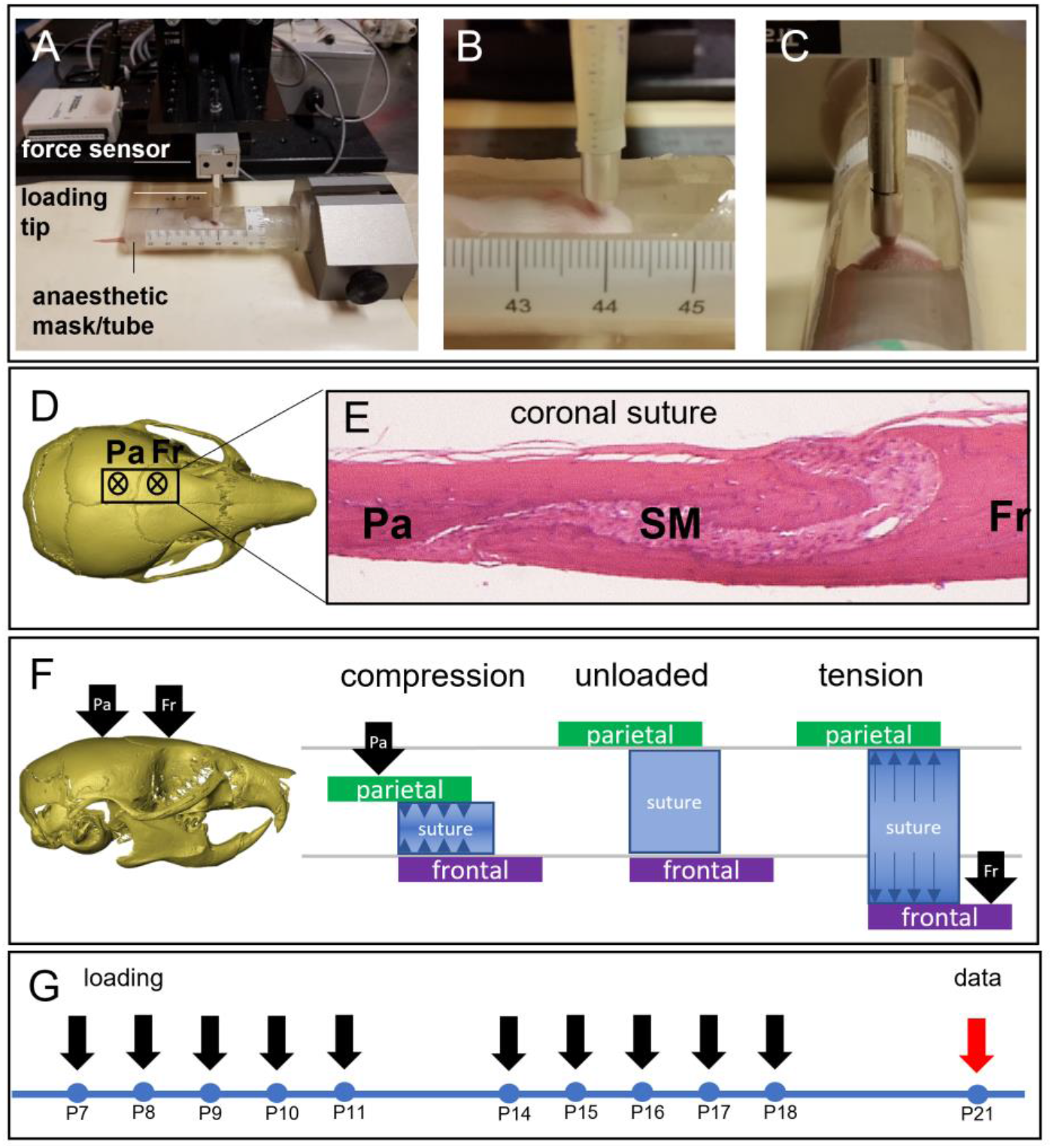
Cranial bone loading on pre-weaning Crouzon mice using actuator technology. **A** Equipment set-up showing P10 mouse in a custom-designed, 3D-printed anaesthetic mask and holder, with a window for access to the skull vault. The electrical, linear actuator moves a small probe (1 mm) that loads the bone underneath, a sensor records the force applied. **B**, **C** Close-ups of the loading protocol in progress at postnatal day 7 (**B**) and 21 (**C**) with the actuator probe targeting the frontal bone. **D** Top view of a micro-CT generated image of a wild-type mouse skull (P21). The asterisks indicate the approximate loading locations on the parietal (Pa) and frontal (Fr) bones, either side of the coronal suture. **E** Histological, sagittal section (H&E) through an adult mouse skull showing the parietal (Pa) and frontal (Fr) bones and the suture mesenchyme (SM) situated between the overlapping parts of the parietal and frontal bones. **F** Schematic representation of the mechanical action generated by loading on either the parietal (Pa) or frontal (Fr) bones. Loading the frontal bone will generate tensile forces on the suture (right) and loading the parietal bone will generate compression forces on the suture (left). **G** Treatment schedule for the cranial loading protocol starting at P7 and loading 10 days until P18. Skulls are collected at P21.

### 3.2. Changes in Crouzon skull morphology after cranial bone loading

We applied cyclical, mechanical loading to the frontal bone of wild-type (WT; n=15) and FGFR2-C342Y (MUT; n=7) mice and to the parietal bone of WT (n=9) and MUT (n=8) mice for 10 days between P7 and P18. On the last day of loading, mice were injected with Alizarin complexone and on P21 mice were culled and skulls dissected. Skulls were then microCT scanned for further morphological characterisation. In all our subsequent analyses, we compared treated mice with untreated control mice, WT (n=12) and MUT (n=16), at P21. In the untreated group (**Figure 2A**), we could clearly see the typical phenotypic appearance of the Crouzon mouse demonstrated by a domed, brachycephalic skull vault and a shortened maxilla which in the most severe cases resulted in malocclusion of the incisors. Superfical analysis of the surface of the skull vault showed complete synostosis of the coronal suture. Analysis of the mutants loaded on the frontal bone showed a distinctive shift to a more normal skull appearance, with reduced doming of the skull and elongated maxillary area (**Figure 2B**). Mice loaded on the parietal bone showed no improvement of Crouzon-related appearance (**Figure 2C**). No apparent effect was noted on WT loaded skulls. To quantify these gross morphological observations, we performed detailed analysis of skull morphologies (i.e. length, width and height) using the microCT images (**Figures 2D-F**). Untreated mutant animals showed a significant difference in the skull width (Δ4.1%, p=1.3E-07), length (Δ12.2%, p=9.2E-11), height (Δ13.4%, p=2.1E-12) when compared to the untreated WTs, represented by the phenotypic effects of midfacial hypoplasia and brachycephaly. Comparing the skull morphologies of the treated and untreated mutant mice, showed distinct differences suggesting phenotypic rescue, but only in the group treated on the frontal bone. Again, no difference was noted between treated and untreated WT skulls. The dimensions of the untreated mutant skull were used as control in all subsequent comparisons to compare the effect of the treatment. Mice treated on the frontal bone showed a statistically significant reduction of the maxillary shortening compared to untreated mutants (skull length: Δ5.9% vs Δ12.2%, p=2.9E-04). Brachycephalic appearance measured by the skull height was reduced in frontal loaded skulls (Δ10.7% vs Δ13.4%, p=0.14), but not statistically significant. Unexpectedly, skull width was significantly different in both the frontal and parietal loaded skulls (Δ7.5% vs Δ4.1%, p=4.7E-04 and Δ7.8% vs Δ4.1%, p=2.9E-04 respectively). Overall, the mutant mice loaded on the frontal bone displayed a restored skull morphology, representing partial phenotypic rescue. In contrast, mice loaded on the parietal bone did not.

**Figure 2.**
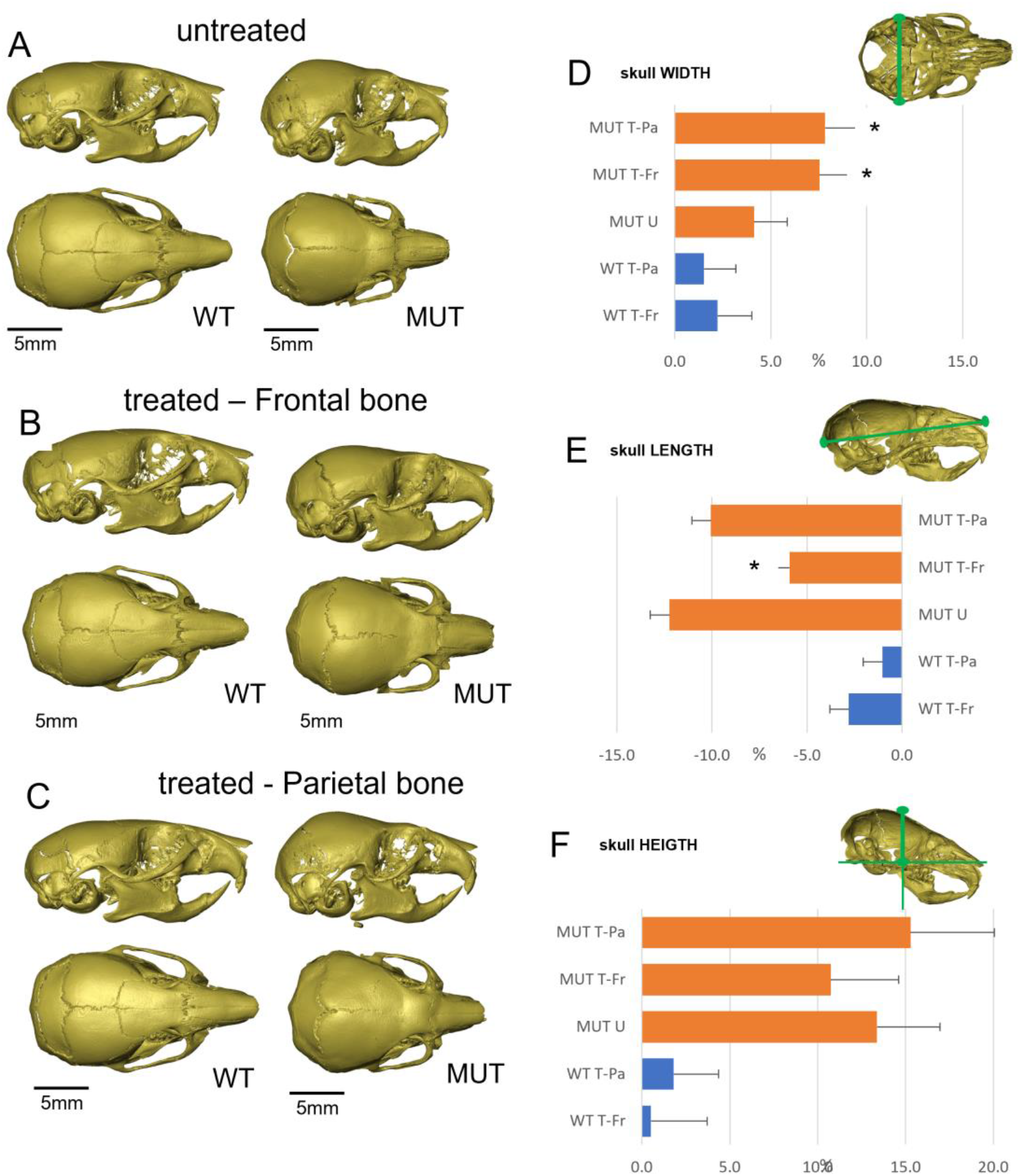
Changes in Crouzon skull morphology after cranial bone loading. **A** micro-CT images of untreated P21 skulls. MUT (Fgfr2_C342Y) skulls show typical signs of brachycephaly and synostosis of the coronal suture. **B** micro-CT images P21 skulls loaded on the frontal bone. While WT skulls appear unchanged, MUT skulls display a partially restored skull shape with a reduced brachycephalic appearance. Synostosis of the coronal suture does not appear to be affected. **C** micro-CT images P21 skulls loaded on the parietal bone. Compared to untreated skulls, no changes can be observed. **D-F** Graphs representing relative skull dimensions in percentage increase/decrease compared to untreated WT controls. **D** Skull measurements show that frontal and parietal loading increases mutant skull width by respectively 7.5% and 7.8% as compared to 4.1% in untreated mutant skulls. **E** Untreated, mutant skulls are 12.2% shorter than untreated WT skulls, indicating the midfacial hypoplasia phenotype. Only frontal bone loaded skulls show a statistically significant difference and are only 5.9% shorter. **F** Untreated, mutant skulls are 13.4% higher than untreated WT skulls, indicating the brachycephalic appearance of these skulls. No statistically significant differences are found for relative height in treated skulls. Images in A-C are representative of the average dimensions per experimental group. Untreated-WT n=12, frontal loaded-WT (WT T-Fr) n=15, parietal loaded-WT (WT T-Pa) n=9, untreated MUT (MUT U) n=16, frontal loaded MUT (MUT T-Fr) n=7, parietal loaded MUT (MUT T-Pa) n=8. *= statistically significant (p<0.05) based on T-test performed against MUT U. Error bars in D-F represent approximate variance calculated using the Delta method.

### 3.3. Changes in coronal suture morphology and histology following cranial bone loading

In an attempt to explain the gross phenotypic effects on skull size and shape after cranial bone loading, we analysed coronal suture morphology in more detail. Using a fluorescent Alizarin Red dye, injected in animals at the end of the loading protocol, we were able to image the calvarial bones and sutures post-treatment at P21. Untreated WT skulls showed the charatcteristic overlapping feature of the coronal suture (**Figure 3A**), while untreated mutants showed loss of suture patency as well as increased bone denisty around the location of the suture (**Figure 3B**). Skulls loaded on the frontal bone showed calvaria with patent sutures in most cases, but with an abnormal suture morphology, including a reduced area of frontal-parietal overlap (**Figure 3C**). Parietal loaded skulls showed no difference from untreated controls (**Figure 3D**). When analysing the patency of the coronal suture in this way we observed that some skulls exhibit an asymmetrical partial rescue of coronal synostosis on the treated (left) side. Other skulls show a wider effect where both sutures remained patent at P21. When using an unbiased (i.e. randomised and blinded) suture patency scoring system, loaded sutures were more likely to be patent or only partially fused than controls. This effect was larger on the treated side (X^2^=9.61, p=0.0019) than on the untreated side (X^2^=4.23, p=0.0397). Histological data of the left coronal suture (**Figure 3E**) confirmed the morphological observations. Untreated WT sutures displayed the typical overlapping structure, with the parietal bone on top of the frontal bone and the sutural mesenchyme in between (**Figure 3F**). Untreated mutant sutures had lost the normal, patent structure and instead showed scattered areas of mesenchyme surrounded by a larger amount of bone (**Figure 3G**). Sutures loaded under tension (frontal bone) had increased patency, but also showed increased bone thickness as seen in untreated sutures (**Figure 3H**). The number of mesenchymal cells in the suture appeared to be decreased compared to WT. Sutures loaded under compression (parietal bone) showed no difference compared to the untreated controls (**Figure 3I**). Using the same histological preparations we analysed the brain tissue underneath the location of loading to exclude any secondary, adverse effects on the underlying cortex. We observed no signs of abnormal cortical lamination (**Supplementary Figure 1**). Therefore, we can conclude that cranial loading on the frontal bone, inducing tension on the suture, causes partial rescue in the majority of cases and total rescue in a minority of cases without any damage to the underlying brain.

**Figure 3.**
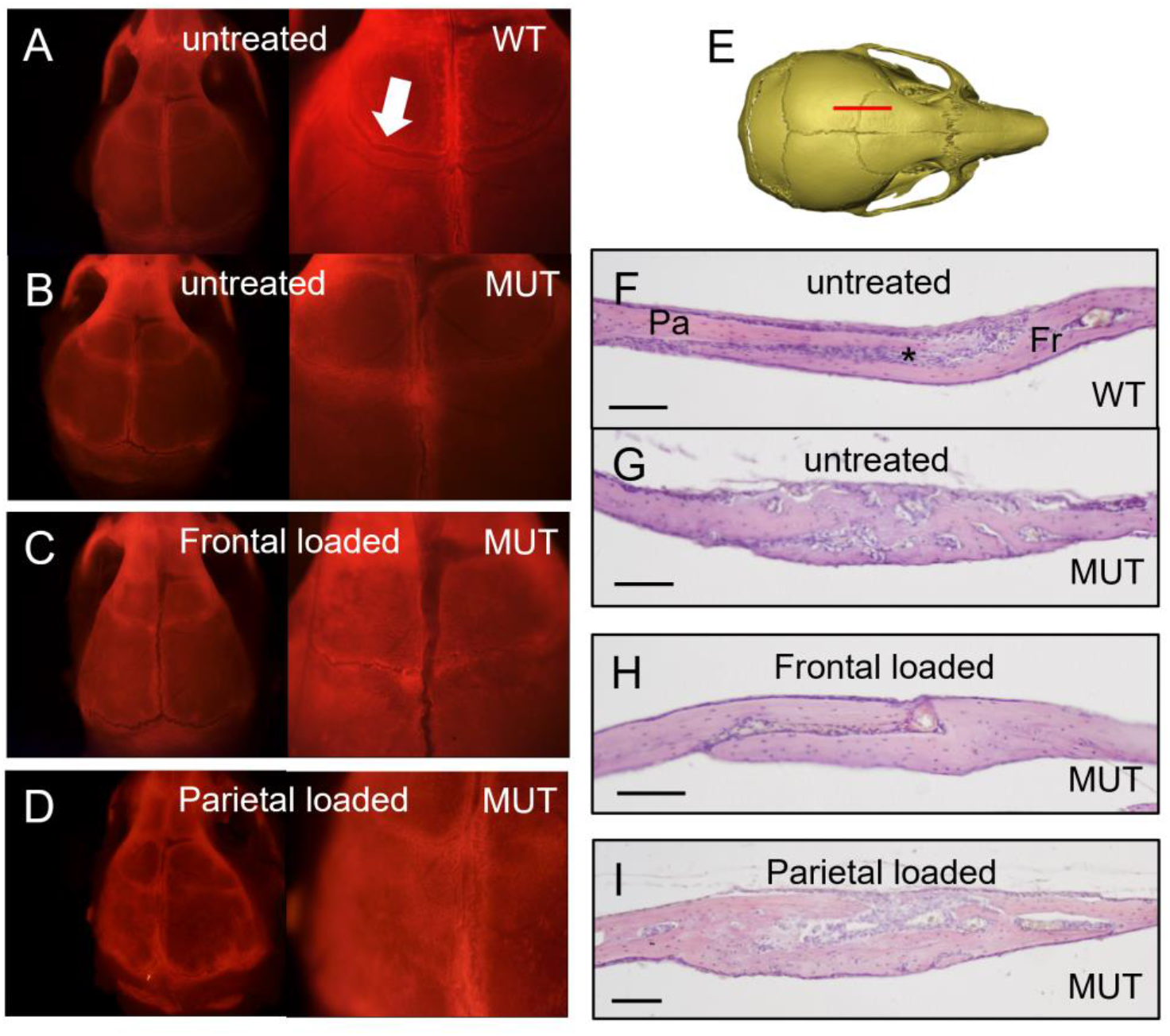
Changes in coronal suture morphology and histology following cranial bone loading. **A-D** Calvaria showing fluorescent Alizarin complexone staining at 0.8x (left) and 2x (right) magnification. **A** Wild-type (WT) calvaria show the typical overlapping bone structure of the coronal suture (arrow). **B** Mutant (MUT) coronal sutures show some staining indicating active bone growth but the overlapping structure is absent. **C** Frontal loaded calvaria show a patent suture, but with a reduced area of frontal-parietal overlap. **D** Parietal loaded calvaria show no difference from untreated mutant sutures. **E** Sagittal sections through the coronal suture (red line) at P21 were stained with H&E. **F** Wild-type (WT) sutures show a characteristic overlapping feature of the parietal (P) and frontal (F) bones with the suture mesenchyme (*) in between. **G** Mutant (MUT) sutures show a loss of patency phenotype with fusion between parietal and frontal bones and loss of mesenchyme. The suture appears thickened as compared to flanking bone. **H** Animals treated with frontal bone loading show a more normal suture with bones appearing unfused, but with significant loss of mesenchyme. **I** Animals treated with parietal bone loading show no improvement and look similar to untreated sutures. Scale bars are 100 μm.

### 3.4. Changes in skull base morphology and histology following cranial bone loading

Analysis of the gross skull morphology showed that mutant mice loaded on the frontal bone display a reduced maxillary shortening (**Figure 2E**). While the delayed synostosis of the coronal suture may contribute to this, we explored whether the cranial base plays a role. Using the micro-CT data, we constructed 2D images of the midline through the cranial base to assess the morphology (**Figure 4A**). At P21, both the sphenoidal-occipital synchondrosis (SOS) and the intersphenoidal synchondrosis (ISS) were open in WT mice, but in the untreated mutants, the ISS is partially or fully closed (**Figure 4B**). This contributes to the maxillary reduction and the resulting midfacial hypoplasia seen in Crouzon mice. The majority of mutant mice loaded on the frontal bone had a partially open ISS, sometimes with a bony bridge located on the caudal side of the cranial base (**Figure 4C**). Parietal bone loaded mutants showed no difference to untreated controls (**Figure 4D**). An unbiased (i.e. randomised and blinded) synchondrosis patency scoring system showed that the increased patency of the ISS in the mutant mice treated on the frontal bone was statistically significant (p<0.05; **Figure 4E**). Histological analysis of the skull base (**Figure 4F**) confirmed that the ISS was abnormal in the mutant mice compared to WT while the SOS was normal in both (**Figure 4G-H**). The mutant ISS cartilage appears ectopic to the cranial base with a reduced number of chondrocytes and is cone-shaped rather than rectangular. Frontal loaded mutant mice show normal ISS cartilage comparable to the untreated WT mice (**Figure 4I**), but parietal loaded mice showed no difference to untreated controls (**Figure 4J**). This data supports an interpretation that cranial loading has an effect on the cranial base, which in turn, contributes to the partially restored skull shape, in particular skull length.

**Figure 4.**
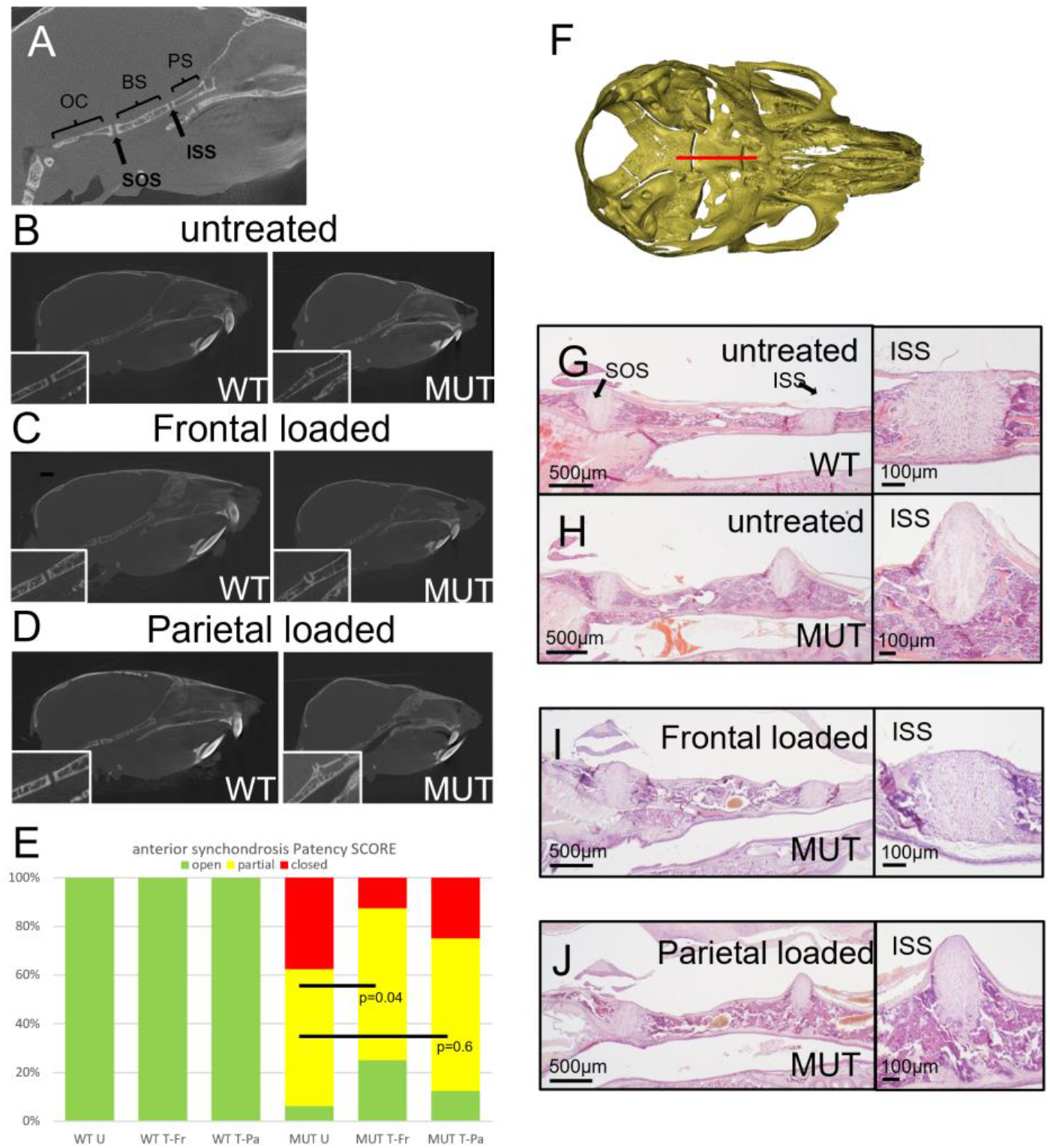
Changes in skull base morphology and histology following cranial bone loading. **A** Schematic of the skull base showing anatomical structures. OC=occipital; BS=basi-sphenoid; PS=pre-sphenoid;SOS=spheno-occipital synchondrosis; ISS=intersphenoidal synchondrosis. **B**-**D** Micro-CT images of sagittal, midline sections through the skull. Untreated WT skulls show a fully patent ISS (inset) while MUT synchondroses have closed (**B**). Frontal loaded skulls show improved patency of the ISS (**C**), while parietal loaded skulls do not (**D**). **E** Patency of the anterior synchondrosis (ISS) was scored blinded for sample identity. While all WT synchondroses are open (green) at P21, the majority of the untreated MUT skull bases have a partially (yellow) or fully closed (red) ISS. Skulls loaded on the frontal bone show a statistically significant improvement while parietal loaded skulls do not. Untreated-WT (WT U) n=12, frontal loaded-WT (WT T-Fr) n=15, parietal loaded-WT (WT T-Pa) n=9, untreated MUT (MUT U) n=16, frontal loaded MUT (MUT T-Fr) n=7, parietal loaded MUT (MUT T-Pa) n=8. **F** Sagittal, midline sections through the skull base (red line) at P21 were stained with H&E. **G-J** Histological analysis of the Crouzon-associated phenotypic features following cranial loading. **G** A wild-type skull base showing the bones seperated by sychondroses. SOS=spheno-occipital synchondrosis; ISS=intersphenoidal synchondrosis. The magnified image of the ISS shows a rectangular cartilage situated between endochodral bone. **H** Mutant (MUT) ISS shows an aberrant cone shape and the cartilage structure protrudes cranially. **I** Animals treated with frontal bone loading show a more normal synchondroses. **J** Animals treated with parietal bone loading show no improvement and look similar to untreated synchondroses.

### 3.5. Comparing the pattern of mechanical strain distribution across the skull between frontal and parietal loading

To understand the pattern of mechanical strain distribution across the skull due to the loading at the frontal and parietal, we used a previously validated finite element (FE) model^25,26^. The FE model was developed based on a P3 WT mouse skull and was used to model calvarial growth. We used this model to predict the shape and pattern of tissue differentiation at P7 and loaded the model at P7 analagous to our *in vivo* experiments (**Figure 5A**). The pattern of first principal strain highlighted that frontal loading induced a lower level of strain across the coronal suture (see highlighted dash boxes in **Figure 5B&C**) compared to parietal loading. It also showed that the load applied to the skull roof induced a mechanical strain on the skull base. Interestingly, the level of induced strain across the intersphenoidal synchondrosis (ISS) was lower in the frontal loading compare to the parietal loading group (see highlighted dash boxes in **Figure 5D&E**).

**Figure 5.**
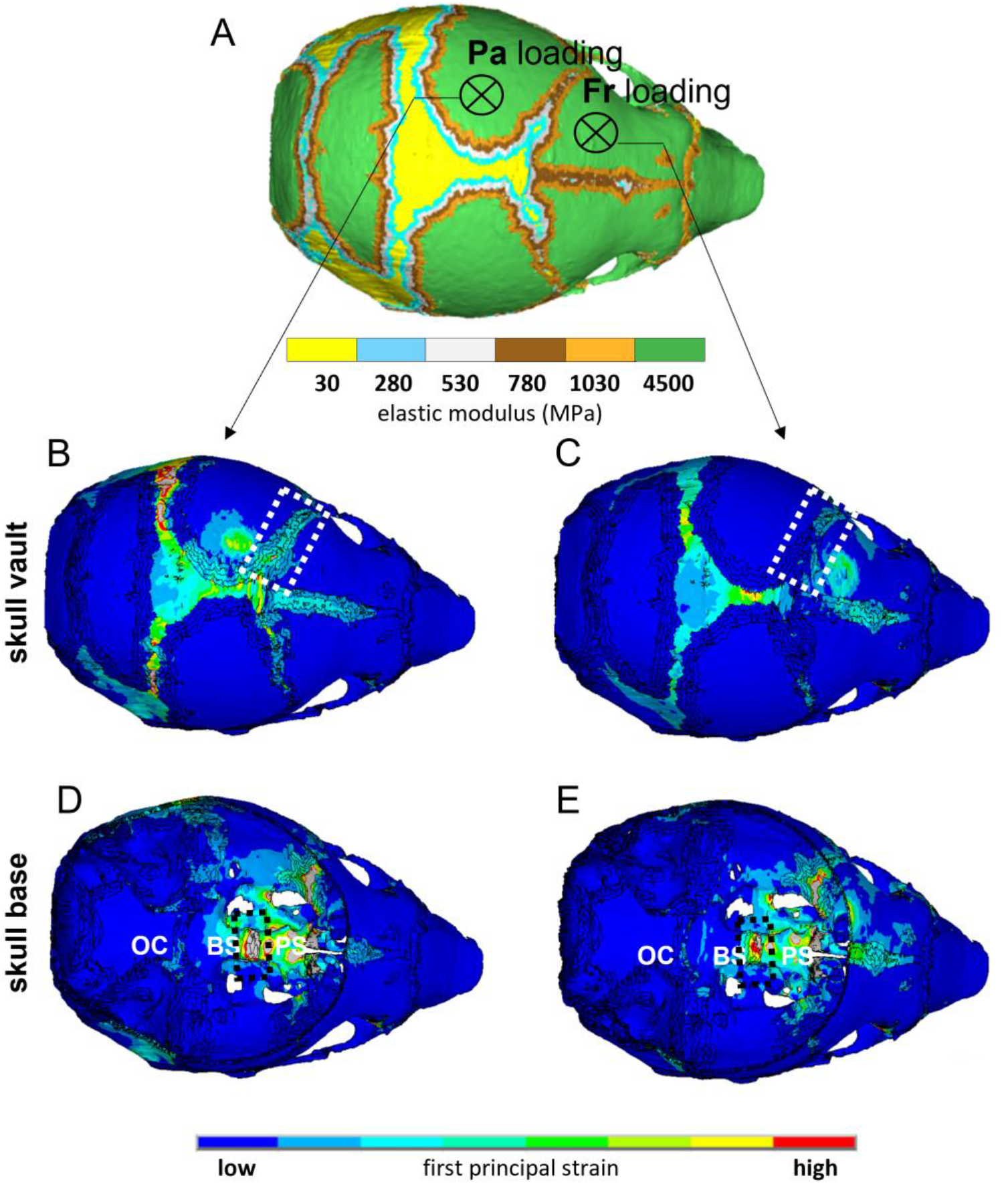
Comparing the pattern of mechanical strain distribution across the skull between frontal and parietal loading. **A** Finite element model of a P3 mouse skull that was computationally grown to P7 and used to investigate the difference in the pattern of strain distribution due to the frontal and parietal loading. It indicates the mechanical properties of different skull regions and the anatomical position that were loaded analogous to the *in vivo* experiments. Heat maps were generated using ANSYS v.18 ([ANSYS Inc](https://www.ansys.com/en-gb)., Canonsburg, PA, USA). **B-C** Comparing the pattern of first principal strain across the skull roof between parietal and frontal loading respectively. The dashed box highlights the coronal suture. **D-E** Comparing the pattern of first principal strain across the skull base between parietal and frontal loading respectivly. The dashed box highlights the ISS (intersphenoidal synchondrosis). OC=occipital; BS=basi-sphenoid; PS=pre-sphenoid.

## 4. Discussion

This study shows that the application of external mechanical loading on the skull of a syndromic craniosynostosis mouse model can prevent the complete closure of the coronal suture and of the anterior synchondrosis, which in turn reduces the damaging effects on skull shape and size, partially restoring normal skull appearance.

Accurate loading of the individual calvarial bones was achieved using a novel set up that was developed enabling us to apply force up to a predetermined maximum of 0.1 N. No adverse effects were observed in any of our loading procedures (total of 670) so we conclude this to be a safe approach for cranial loading in mouse. While the mice were subjected to general anaesthesia, this was mainly to avoid any movement during the procedure and we don’t foresee this to be necessary when applied to patients. Further work will focus on the optimisation of the force/frequency needed to achieve the desired effect and on answering the question whether an initial loading treatment is sufficient to prevent complete synostosis or if loading needs to be applied long term to avoid the process of synostosis recommencing. Preliminary investigations showed that the cortical lamination of the brain remains intact after loading treatment, but further behavourial studies will be needed to confirm this.

Overall we identified a significant effect on the size and shape of Crouzon skulls after frontal loading only. This can be explained by the characteristic feature of the coronal suture, where the parietal bone partially overlaps the frontal bone. Loading the frontal bone will load the soft sutural tissue under tension, that is likely to cause a certain level of shear stress. Cellular tension has been shown to activate the ROCK/TAZ pathway in vitro^28^, but further *in vivo* analysis of the mechanotransduction effects on the sutural mesenchyme can potentially unravel the precise mechanism. While we anticipated the effect to be more pronounced on the loaded side (L) compared to the unloaded side (R), analysis showed only a small number of samples that presented with an asymmetric presentation of coronal synostosis rescue. A likely explanantion for this is that the mechanical strain that has been induced as a result of the unilateral loading transfers onto the unloaded coronal suture on the opposite side. The finite element data confirm that a low level of strain was indeed induced on the unloaded coronal suture albeit higher in the parietal loaded group compare to the frontal loaded group. At the same time, our observation that cranial loading prevents complete fusion of the synchondrosis supports the theory that loading the calvarial can impact the skull base through the 3D structural framework of the cranium.

In our hands, the Crouzon Fgfr2-C342Y mouse model on a CD-1 background shows near complete fusion of the coronal suture at P21, the end point of our experiments^20,24^. Compared to unloaded controls, frontal loaded sutures showed a mixed histological picture with increased patency on the one hand and reduction of the sutural mesenchyme on the other hand. Disappearance of the sutural mesenchyme by premature differentiation into osteoblast and bone is typical of the pathological synostosis process caused by FGFR2 mutation^29^ and allows the fusion of the flanking bones. In this study we show that the daily application of mechanical tension prevents the fusion, while the reduction in sutural mesenchyme suggests that it is unable to prevent the premature differentiation of the mesenchyme. As a result, the coronal suture retains it’s patency allowing the skull to grow normally without the compensatory growth associated with coronal synostosis^30^.

An additional, unexpected discovery was that apart from partially rescueing the coronal suture synostosis phenotype, we were also able to partially rescue the premature closure of the anterior synchondrosis (i.e. intersphenoidal synchondrosis (ISS)) in the skull base of the Crouzon mouse. Prevention of premature fusion of the ISS in this mouse model for syndromic craniosynostosis is likely to contribute to the normalisation of skull length that was observed in this study. Anatomically, the mouse ISS is similar to that in human and natural fusion of the human ISS occurs between 2-4 years of age^31^. There is no CT data available for the premature fusion of the ISS in human Crouzon syndrome patients, but in the Crouzon mouse model the ISS is fused prematurely at P21 as shown by by Liu et al.^23^ and this study. A significant delay in the premature fusion was identified in the frontal loaded Crouzon mice, which suggests that cranial loading transfers its effect indirectly to the synchondroses in the skull base. This was also corroborated by our finite element results that showed that calvarial loading induced mechanical strain on the skull base and ISS. Histologically, the mutant ISS displays as a cone shape cartilage, protruding rostrally from the sphenoidal flanking bones with a bony bridge forming caudally. We speculate that applying mechanical tension on the anterior side of the ISS (i.e. the presphenoidal bone) causes shear stress on the caudal side preventing the formation of the bony bridge and the subsequent protrusion of the cartilaginous synchondrosis. Alternatively, a mechanotransductional effect on the developing chondrocytes in the synchondrosis is possible, as it is known that the formation and fusion of the ISS is regulated by the Indian and Sonic Hedgehog pathways^32^. Clinically, skull base abnormalities are difficult to resolve surgically because of the anatomical position of the target tissue. External application of a corrective mechanical force may provide be a possible way to address this clinical challenge.

It is clear that the proposed approach in this study can be further optimised and needs to undergo a feasibility study in a larger animal model in preparation for clinical translation if its efficacy is proven. At the same time, expanding this treatment approach to other transgenic mouse models of syndromic craniosynostosis (e.g. Apert syndrome) will clarify if mechanical loading will be universally effective on other types of craniosynostosis, especially other types of sutures. At present it seems that a mechnical loading approach can only be deployed clinically on patients that present with sutures that are not fully fused i.e. have not completed the synostosis process. The number of craniosynostosis patients that present with partially patent sutures is estimated to be 5-10% at Great Ormond Street Hospital (D Dunaway 2021, personal communiation). Another potential application of the cranial loading can be to prevent the healing of surgically seperated calvarial bones post-operatively (aka resynostosis). This is an important clinical problem causing significant morbidity leading to the need for repeated reoperations. Attempts to prevent reoperation have been made using glypican-3 containing nanotubes to inhibit cranial bone regeneration in a Crouzon mouse model and by regenerating a functional suture using Gli+ mesenchymal stem cells in a Twist1 mouse model of craniosynostosis^33,34^. While these potential treatments have not been applied to patients yet, they will be invasive procedures involving the introduction of pharmacological or cellular agents with potential unwanted side effects. It may be possible to achieve a similar effect using cranial loading on patients post-surgery.

## 5. Conclusion

In summary, *in vivo* calvarial loading using pre-weaning Crouzon mice shows a distinct phenotypic effect indicated by a partially restored skull morphology and improved patency of the coronal suture and anterior synchondrosis. Further studies into the use of mechanotransduction to delay or prevent premature, pathogenic suture closure have the potential to be a first important step towards the development of a non-invasive, non-surgical treatment alternative for children with progressive craniosynostosis.

## Acknowledgements

The *Fgfr2c*^C342Y^ mouse colony was derived from the EMMA Consortium. This work was supported by Newlife, the charity for disabled children (award 17-18/18), the Royal Academy of Engineering (10216/119); and the Engineering and Physical Science Research Council (EP/W008092/1) and University College London. EP is a GOSHCC Principal Investigator. This research was supported by the NIHR Great Ormond Street Hospital Biomedical Research Centre. The views expressed are those of the author(s) and not necessarily those of the NHS, the NIHR or the Department of Health. We are grateful to Michael Fagan for micro-CT scanning support and to Susan Herring for advice and support throughout this study.

## Author contributions

M.M., E.P. and A.A. conceived and designed the project; M.H., D.S., M.M., A.M., D.J. and E.P. acquired, analysed and interpreted data; E.P. and M.M. wrote the paper; all authors commented on and contributed to the paper.

## Data availability

The datasets generated during and/or analysed during the current study are available from the corresponding author on reasonable request.

## Supplementary figures

**Supplementary Figure 1.**
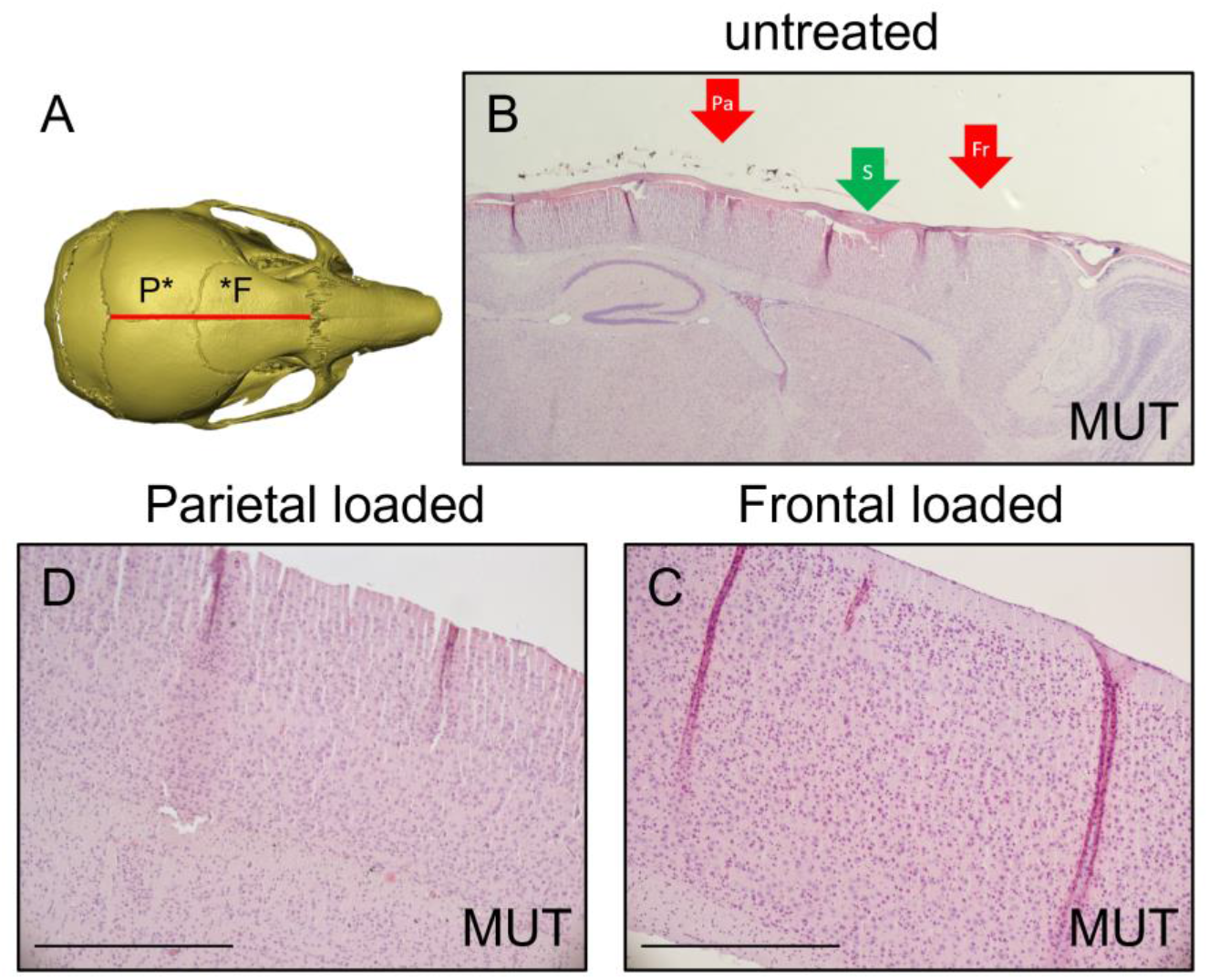
Histological analysis of brain tissue following cranial bone loading. **A** Sagittal, midline sections through the brain (red line) at P21 were stained with H&E. **B** Low magnification image of a mutant brain showing the fused coronal suture (green arrow, S) as well as the approximate location of frontal (red arrow, Fr) and parietal (red arrow, Pa) loading. **C** High magnification image of a frontal loaded mutant brain showing normal cortical lamination. **D** High magnification image of a parietal loaded mutant brain showing normal cortical lamination.

**Supplementary Video 1a** Video footage (25s) of cranial bone loading treatment protocol. This video shows a P7 WT mouse being loaded on the frontal bone.

**Supplementary Video 1b** Video footage (25s) of cranial bone loading treatment protocol. This video shows a P14 WT mouse being loaded on the frontal bone.

**Supplementary Video 2a** Video footage (52s) showing behaviour of three WT P21 litter mates (L19, L20 and L21) after completing the two-week cranial bone loading treatment protocol.

**Supplementary Video 2b** Video footage showing behaviour of three MUT P21 litter mates (L18, L25 and L26) after completing the two-week cranial bone loading treatment protocol.

